# Microbiome imaging goes à la carte: Incorporating click chemistry into the fluorescence-activating and absorption-shifting tag (FAST) imaging platform

**DOI:** 10.1101/2023.10.02.560575

**Authors:** David M. Anderson, Matthew G. Logan, Sara S. Patty, Alexander J. Kendall, Christina Z. Borland, Carmem S. Pfeifer, Jens Kreth, Justin L. Merritt

## Abstract

The human microbiome is predominantly composed of facultative and obligate anaerobic bacteria that live in hypoxic/anoxic polymicrobial biofilm communities. Given the oxidative sensitivity of large fractions of the human microbiota, green fluorescent protein (GFP) and related genetically-encoded fluorophores only offer limited utility for live cell imaging due the oxygen requirement for chromophore maturation. Consequently, new fluorescent imaging modalities are needed to study polymicrobial interactions and microbiome-host interactions within anaerobic environments. The fluorescence-activating and absorption shifting tag (FAST) is a rapidly developing genetically-encoded fluorescent imaging technology that exhibits tremendous potential to address this need. In the FAST system, fluorescence only occurs when the FAST protein is complexed with one of a suite of cognate small molecule fluorogens. To expand the utility of FAST imaging, we sought to develop a modular platform (Click-FAST) to democratize fluorogen engineering for personalized use cases. Using Click-FAST, investigators can quickly and affordably sample a vast chemical space of compounds, potentially imparting a broad range of desired functionalities to the parental fluorogen. In this work, we demonstrate the utility of the Click-FAST platform using a novel fluorogen, ^PL^Blaze-alkyne, which incorporates the widely available small molecule ethylvanillin as the hydroxybenzylidine head group. Different azido reagents were clicked onto ^PL^Blaze-alkyne and shown to impart useful characteristics to the fluorogen, such as selective bacterial labeling in mixed populations as well as fluorescent signal enhancement. Conjugation of an 80 Å PEG molecule to ^PL^Blaze-alkyne illustrates the broad size range of functional fluorogen chimeras that can be employed. This PEGylated fluorogen also functions as an exquisitely selective membrane permeability marker capable of outperforming propidium iodide as a fluorescent marker of cell viability.

## Main

Recent estimates for the number of resident bacteria residing on and within a typical human host average about 3.8 x 10^13^ cells, a roughly one-to-one ratio with the total number of host cells^1^. The vast majority of this population would be classified as anaerobic in nature, either requiring strictly anoxic (<0.5% oxygen) or microaerophilic (∼2-8% oxygen) conditions. Anaerobic organisms in the oral cavity^2^ and lower intestine^3^ outnumber their aerobic counterparts by 2-3 orders of magnitude; even the human skin microbiota is dominated by anaerobes^4^. The microbial ecology of these anaerobic mucosal communities is a central determinant of mucosal health and is thus a major focus of research in the field. Disruptions to this ecology (referred to as dysbiosis) can result in a wide array of chronic inflammatory diseases of the mucosae. Mucosal dysbiosis is also increasingly recognized for its systemic health effects as well. For example, the oral microbiota and its byproducts have been associated with a wide range of extraoral conditions, including: Alzheimer’s disease, inflammatory bowel disease, diabetes, arthritis, atherosclerosis, premature birth, and head/neck, gastric, pancreatic, and colorectal cancers ^5^.

The anoxic growth conditions needed to study the biological mechanisms of these organisms poses a unique challenge for fluorescence microscopy studies due to the strict oxygen requirement for the most commonly employed genetically-encoded fluorescent proteins like the green fluorescent protein (GFP) and others. Thus, there has been a longstanding need in the field to develop new genetically-encoded anaerobic live-cell imaging tools. One promising method to address this gap employs a small (14 kDa) monomeric protein known as the fluorescence-activating and absorption-shifting tag (FAST). The natural scaffold to this protein originated from the yellow protein isolated from *Halorhodospira halophila* by T.E. Meyer ^6^. Plamont and coworkers have since developed the first iteration of the current FAST variants through directed evolution ^7^. That work included the origination of the FAST ligand, a hydroxybenzylidine-rhodanine conjugate whose fluorescence requires complexation with the FAST protein. Subsequent versions of FAST proteins have since been constructed, including: biochemically-improved ^8^, red-shifted ^9^, split ^10^, promiscuous ^11^, nano-sized ^12^, and chemically-assisted tethering^13^ versions. Further exciting advances in companion fluorogen chemistries beyond the original 4-hydroxybenzylidene-rhodanine (HBR) or 4-hydroxy-3-methylbenzylidene-rhodanine (HMBR) scaffolds have also been achieved ^7^, including the introduction of a far-red emitting FAST fluorogen ^9^, GFP-chromophore mimetics ^14,15^, and stimuli-responsive derivatives ^16^.

Building upon these advancements, we sought to exploit the unique characteristics of FAST fluorescence to develop a new concept in fluorescent imaging: user-defined modularity. To achieve this, we synthesized two FAST fluorogen variants capable of harnessing the power of click chemistry to support personalized, on-demand fluorogen modifications for unique biological applications. One of these is a new fluorogen derived from the extremely inexpensive reagent ethylvanillin^17^. Copper catalyzed click chemistry^18,19^ was used to conjugate several commercially available azido reagents to this fluorogen via its rhodanine-alkyne appendage to illustrate the potential utility of Click-FAST for a diverse range of biological applications. The simplicity, specificity, and exceptional efficiency of the click reaction provides Click-FAST users with unique access to the benefits of synthetic chemistry without requiring specialized knowledge or equipment to develop new fluorogen chemistries and bioactivities for a near limitless range of fluorescence imaging applications.

## Results

### The FAST system excels in an anaerobic environment

Oxygen concentration is a critical factor modulating the photostability of fluorescent proteins during imaging, as free singlet oxygen and oxygen radicals can catalyze damaging reactions to the protein chromophores ^20^, while also quickly reducing the viability of oxygen-sensitive anaerobes. Unlike GFP and related proteins, the FAST system has been demonstrated to function normally in anaerobic conditions^21^, but to our knowledge it has not been comparatively demonstrated how gradients of oxygen impact the performance of FAST. Thus, we directly captured fluorescence emission in a regulated environmental chamber to assess photobleaching kinetics. These experiments employed a condon-optimized version of FAST constitutively expressed from a shuttle vector in the oral pathobiont *Streptococcus mutans* following exposure to a cocktail of translation-inhibiting antibiotics. Using a minimal supernatant volume containing the fluorogen ^TF^Coral ^22^, we could measure a distinct gradient of photobleaching rates that directly correlated with oxygen concentration (**Fig. 1a**). At 0.1% oxygen, we observed exceptional photostability, with minimal photobleaching detectable after ten minutes of wide-field illumination. This is in stark contrast to the results observed with ambient levels of atmospheric oxygen, which caused fluorescence to drop precipitously within minutes, similar to what would normally be observed with GFP and related proteins. A final acquisition series performed after equilibrating the chamber back to 0.1% oxygen demonstrated a reversion to high photostability, providing further evidence for the role of oxygen in fluorescence photobleaching. The recovery in signal suggests that the bleaching observed was fluorophore dependent, as translation inhibition would have prevented new synthesis of FAST proteins. Given the extraordinary anaerobic photostability of FAST fluorescence, one could theoretically image a broad range of FAST reporter constructs, including those with relatively weak fluorescence.

**Fig. 1.**
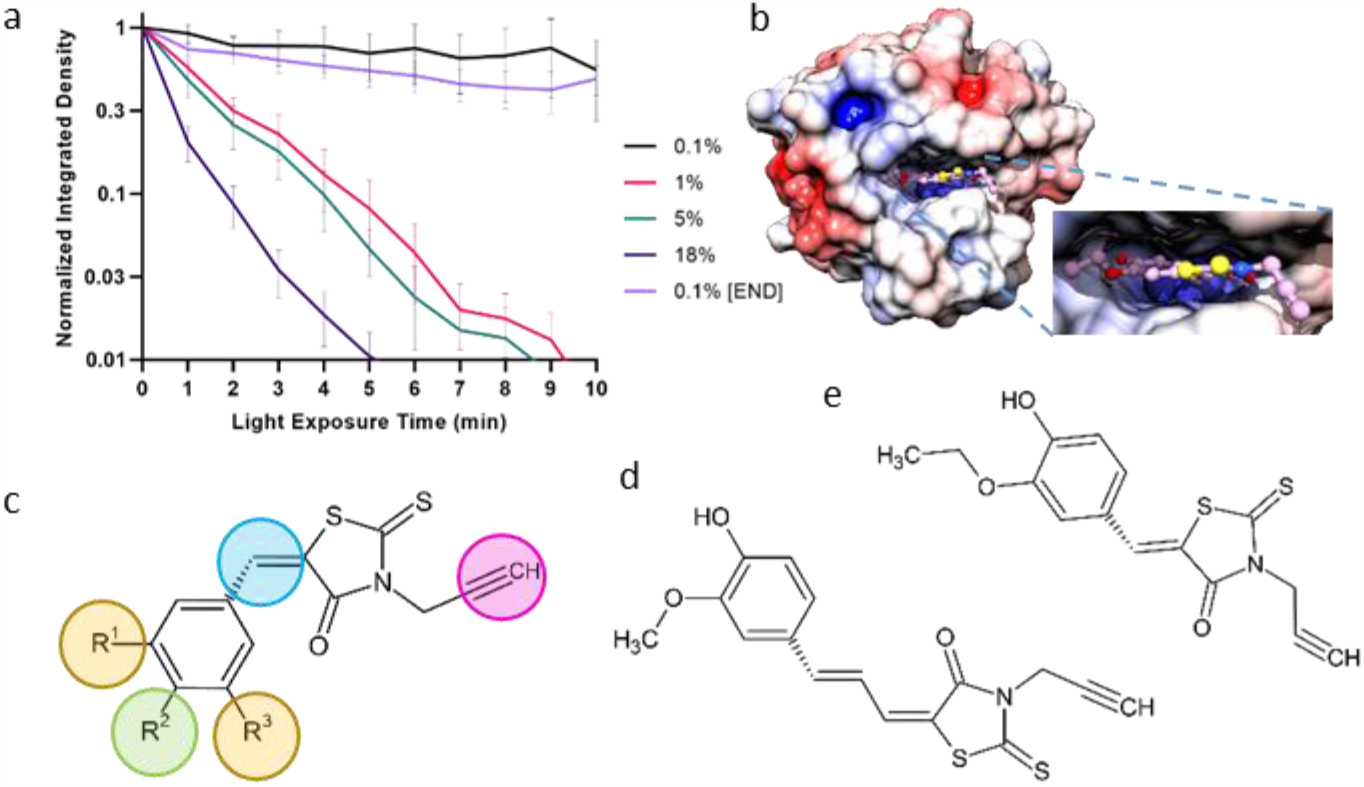
Rationale for clickable FAST ligand. **a**, Stability profile of ^TF^Coral signal over several minutes of exposure in the indicated percentage of oxygen. The “0.1% END” data was obtained after the 18% oxygen data was acquired. N = 4, ± SEM. **b**, Model of FAST (PDB: 7AVA) with a clickable ^PL^Blaze fluorogen docked into the structure using the S1Q ligand as a guide. Surface charge was calculated using APBS^32^ and is shown on a scale of −5 to 5 k_B_T at pH 7.2 (red, negative; blue, positive). The image was generated using UCSF Chimera^33^. **c**, Schematic of a generalized alkyne-FAST fluorogen. Excitation and emission spectra can be selected by altering a combination of the R^1^/R^3^ side groups (yellow) or by a bathochromatic shift via an extended carbon linker (blue). *In situ* functionality can be altered by modifications to R^2^ (green). The location of the clickable component is shown here as an alkyne (magenta). **d**-**e**, Schematics showing the structures of ^TF^Poppy-Click and ^PL^Blaze-Click fluorogens, respectively.

### Introduction to the Click-FAST concept

Encouraged by these results, we sought to broaden the utility of FAST imaging by developing a simple approach to expand the chemical space of the FAST fluorogen concept utilizing robust yet accessible organic synthesis routes based on click chemistry strategies^23^. A FAST-ligand model based on the structure reported by Mineev^12^ reveals how the fluorogen rhodanine group protrudes out of the FAST protein benzaldehyde binding pocket to a solvent accessible area, representing an attractive site for further conjugation (**Fig. 1b**). When paired with the ever-expanding palette of newly reported FAST fluorogens, the further incorporation of clickable modifications presents an exciting opportunity to develop novel bioactive fluorogens tailor-made for individual use cases (**Fig. 1c**). To test the click concept, two FAST analog fluorogens were synthesized with a clickable alkyne group conjugated to the rhodanine nitrogen atom. One of these utilized the ^TF^Poppy farred fluorogen framework ^9^, while the other is novel to this work, incorporating ethylvanillin as an inexpensive and widely available hydroxybenzylidine head group (**Fig. 1d, 1e, Supplemental Figures 1-4**). This new variant was named ^PL^Blaze due to its light-orange fluorescence emission (575 nm).

### In vitro comparisons of FAST fluorogen alkyne variants

Fluorogen affinity measurements and brightness were queried *in vitro* to assess the impact of click conjugation on their performance with the improved FAST protein (iFAST2) and the far-red FAST (frFAST2) protein variant harboring additional point mutations (**Table 1, Suppl. Fig. 5**). Similar to previous measurements made by others, we observed a submicromolar affinity of both FAST protein variants for ^TF^Coral, as well as for frFAST2 with ^TF^Poppy. As expected, ^TF^Coral also produced exceptionally bright fluorescence, yielding more than twice the maximum brightness (here referred to as F_max_) of ^TF^Poppy bound to frFAST2. The affinity of the iFAST2 protein for ^TF^Poppy was approximately 18-fold lower than that of frFAST2. Our new fluorogen variant ^TF^Poppy-alkyne maintained its selective high binding affinity with the frFAST2 protein. However, its apparent F_max_ exhibited a 30-fold decrease. ^TF^Poppy-alkyne displayed a negligible binding affinity to iFAST2, yielding little to no detectable fluorescence signal above background. In stark contrast to ^TF^Poppy and ^TF^Poppy-alkyne, ^PL^Blaze-alkyne exhibited similarly high binding affinities to both iFAST2 and frFAST2 proteins, yielding affinities of approximately 2.6 and 1.2 micromolar, respectively. As such, ^PL^Blaze-alkyne also yielded strong fluorescence with both proteins, producing approximately 75 and 60 percent of the maximum brightness of ^TF^Coral.

**Table 1.** Apparent affinity and brightness metrics of several FAST fluorogens and variants with IFAST2 or frFAST2.

| Fluorogen | Protein | K<sub>d</sub><sub>app</sub> (μM)<sup>1</sup> | F<sub>max</sub> (RFU × 10<sup>5</sup>)<sup>1</sup> | F<sub>max</sub> / K<sub>d</sub><sub>app</sub> |
| --- | --- | --- | --- | --- |
| <sup>TF</sup>Coral | iFAST2 | 0.53 (0.01) | 2.62 (0.021) | 4.962 |
|  | far-red FAST2 | 0.78 (0.01) | 2.57 (0.029) | 3.295 |
| <sup>TF</sup>Poppy | iFAST2 | 12.90 (3.84) | 0.04 (0.002) | 0.003 |
|  | far-red FAST2 | 0.72 (0.01) | 1.10 (0.003) | 1.528 |
| <sup>TF</sup>Poppy-alkyne | iFAST2 | ND | < 0.001 | ND |
|  | far-red FAST2 | 0.67 (0.01) | 0.03 (0.001) | 0.045 |
| <sup>TF</sup>Poppy-lactose | iFAST2 | 12.20 (6.86) | 0.04 (0.001) | 0.003 |
|  | far-red FAST2 | 6.49 (2.06) | 0.07 (0.003) | 0.011 |
| <sup>TF</sup>Poppy-benzoic acid | iFAST2 | ND | < 0.001 | ND |
|  | far-red FAST2 | 0.25 (0.001) | 0.06 (0.007) | 0.240 |
| <sup>PL</sup>Blaze-alkyne | iFAST2 | 2.67 (0.10) | 1.93 (0.001) | 0.723 |
|  | far-red FAST2 | 1.17 (0.03) | 1.53 (0.001) | 1.308 |
| <sup>PL</sup>Blaze-lactose | iFAST2 | 3.94 (1.01) | 1.62 (0.001) | 0.411 |
|  | far-red FAST2 | 1.82 (0.24) | 1.47 (0.01) | 0.808 |
| <sup>PL</sup>Blaze-benzoic acid | iFAST2 | 1.11 (0.06) | 3.24 (0.02) | 2.919 |
|  | far-red FAST2 | 0.41 (0.01) | 2.45 (0.003) | 5.976 |
| <sup>PL</sup>Blaze-PEG<sup>2</sup> | iFAST2 | 4.05 (1.24) | 0.93 (0.005) | 0.23 |
|  | far-red FAST2 | 2.96 (0.30) | 0.97 (0.001) | 0.33 |

<sup>1</sup>Parentheses indicate the standard deviation of three replicate experiments.
<sup>2</sup>PEG abbreviated from N-(acid-PEG10)-N-bis(PEG10)

### Application of Click-FAST for differential fluorescent labeling of bacteria

We surmised that one useful application of the Click-FAST approach would be to screen for fluorogen modifiers supporting selective labeling of mixed bacteria. For instance, an ideal conjugant would be selectively recognized and imported by a specific strain or cell type while being excluded from others. As a proof of principle, mixed cultures of *Streptococcus mutans* UA159 and *Escherichia coli* BL21 were employed to test the efficacy of fluorogen variant fluorescence in bacteria having both Gram-positive monodermic and Gram-negative didermic membrane architectures. Each of these strains are well-characterized, facultative aerobes with defined expression plasmids we use for this work. Additionally, we make use of a robust fusion system for the extracellular display of FAST via attachment to the lipoprotein-outer membrane protein A (Lpp-OmpA) sequence defined by Earhart^24^. As an initial test for selective fluorogen labeling, we clicked azido-lactose and azido-benzoic acid to both ^TF^Poppy-alkyne and ^PL^Blaze-alkyne to compare their potential utilities. Firstly, we compared the effect of click labeling upon the performance of the fluorogens relative to their parental alkyne molecules. The ^TF^Poppy-lactose conjugate exhibited a 17-fold reduction in affinity for the frFAST2 protein, resulting in ∼10-fold lower theoretical F_max_. Interestingly, ^TF^Poppy-lactose produced measurable binding to iFAST2, whereas the parental fluorogen ^TF^Poppy-alkyne did not. Its affinity for iFAST2 and maximal fluorescence metrics also mirrored the original ^TF^Poppy fluorogen. ^TF^Poppy-benzoic acid retained its selective affinity to frFAST2, effectively doubling the apparent binding affinity and F_max_. There was negligible binding and fluorescence detected between ^TF^Poppy-benzoic acid and iFAST2. Unlike ^TF^Poppy-alkyne, conjugation of lactose to ^PL^Blaze yielded only a slightly reduced FAST protein binding affinity, resulting in strong fluorescence with only modest reductions in F_max_. Conjugation to benzoic acid also increased the apparent affinity of ^PL^Blaze for both iFAST2 and frFAST2, which resulted in a concomitant brightness value 60% greater than the parental fluorogen. A chemical mechanism supporting an increased fluorescence output beyond an increased occupancy of the ligand-binding pocket is not yet clear. Minimal spectral shifting was observed following benzoic acid conjugation and both fluorogens share similar HOMO/LUMO electronic structures (**Suppl. Fig. 6**). Overall, ^PL^Blaze was far more permissive of conjugation to both lactose and benzoic acid compared to ^TF^Poppy.

Since frFAST2 performed well with the clicked conjugates of both ^TF^Poppy and ^PL^Blaze (**Table 1**), these fluorogens were each tested in mixed cultures of frFAST2-expressing *S. mutans* and *E. coli* to compare their performance *in vivo*. Both ^TF^Poppy and ^PL^Blaze conjugated to 4-azidobenzoic acid yielded easily discernable fluorescence from individual cocci within chains of *S. mutans*, whereas fluorescence was virtually undetectable from the rod-shaped *E. coli* (**Fig 2a, 2c**.). Benzoic acid, a close derivative of 4-azidobenzoic acid, is commonly employed as a food preservative and has a minimum inhibitory concentration (MIC) in both organisms that is more than three orders of magnitude higher than what was used for these imaging experiments^25,26^. Thus, it is highly unlikely that toxicity, or a related mechanism compromising the membrane barrier, is responsible for the labeling of *S. mutans*.

**Fig. 2.**
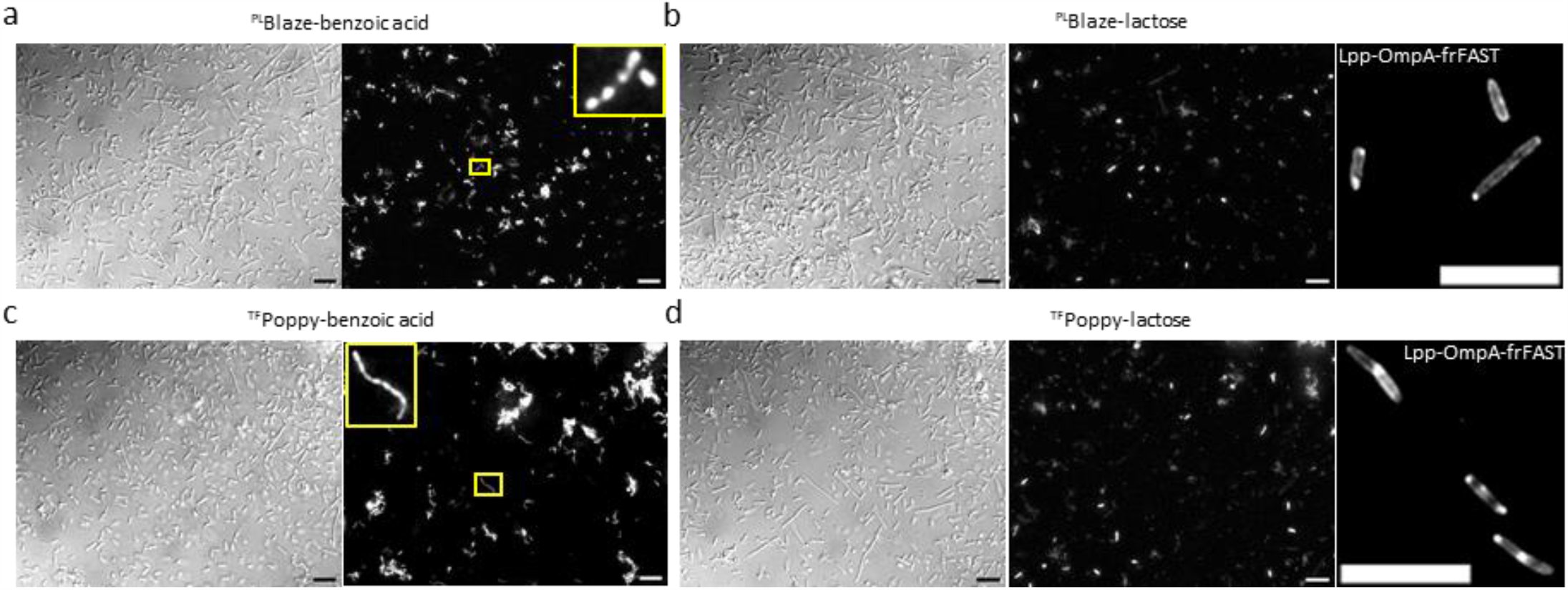
Examples of differential fluorescence signal profiles by clickable FAST fluorogens. **a**, DIC image (*left*) and fluorescence maximum intensity projection image (*right*) of a mixed population of frFAST2-expressing *S. mutans* UA159 and *E. coli* BL21 in the presence of ^PL^Blaze-benzoic acid fluorogen. *Inset*, zoomed image showing labeling of individual *S. mutans* cells. **b**, DIC image (*left*) and fluorescence maximum intensity projection image (*center*) of a mixed population of frFAST2-expressing *S. mutans* UA159 and *E. coli* BL21 in the presence of ^PL^Blaze-lactose. *Right*, Fluorescent image of Lpp-OmpA-frFAST *E. coli* BL21 in the presence of ^PL^Blaze-lactose. **c**, DIC image (*left*) and fluorescence maximum intensity projection image (*right*) of a mixed population of frFAST2-expressing *S. mutans* UA159 and *E. coli* BL21 in the presence of ^TF^Poppy-benzoic acid. *Inset*, zoomed image showing labeling of individual *S. mutans* cells. **d**, DIC image (*left*) and fluorescence maximum intensity projection image (*center*) of a mixed population of frFAST2-expressing *S. mutans* UA159 and *E. coli* BL21 in the presence of ^TF^Poppy-lactose. *Right*, Fluorescent image of Lpp-OmpA-frFAST *E. coli* BL21 in the presence of ^TF^Poppy-lactose. Scale bars = 10 µm.

Click-FAST could also potentially be used to study the import of certain metabolite mimetics into a cell. Thus, we were curious whether a conjugated lactose fluorogen could be imported into these cells. The structure of the *E. coli* LacY permease has been extensively characterized, and analysis of a crystallographically-derived structure of LacY bound to both β-D-galactopyranosyl-1-thio-β-D-galactopyranoside and n-Nonyl-β-D-galactopyranoside suggested to us that it may be feasible to transport a similarly-sized lactose-conjugated fluorogen ^27^. Overall, we observed very weak ^PL^Blaze-lactose fluorescence from *S. mutans* and very bright fluorescence from a small subpopulation of *E. coli* cells, as shown by rodshaped fluorescence. To demonstrate that this effect was not due to impairments in FAST fluorescence, an outer membrane protein A ^13^ chimera with frFAST was constructed, which specifically localizes frFAST to the *E. coli* outer membrane ^24^. When ^PL^Blaze-lactose was added to *E. coli* cells expressing this construct, membrane and polar-localized fluorescence was readily discernable (**Fig. 2b, 2d**). Together, these data demonstrate that lactose-modified fluorogens are excluded from the cytosol in all but a subset of actively growing mid-logarithmic phase *E. coli*, suggesting certain modifications to Click-FAST fluorogens may have utility as tools to reveal subpopulation-specific phenotypes.

### PEG-FAST: a stringently membrane impermeable FAST derivative

Live/dead cellular staining is a standard microscopic approach used to visualize cell viability in bacterial biofilm communities. Nucleic acid-binding dyes such as propidium iodide (PI) are most often employed with the assumption that viable cells specifically exclude these dyes from the cytosol due to membrane impermeability. In practice, we have found that bacteria undergoing periods of starvation and/or stress often result in bright staining with these dyes while the cells are still perfectly viable, especially within biofilms^28^. The FAST-ligand model presented in **Figure 1b** indicates the feasibility of conjugating exceptionally large molecules to FAST fluorogens without impeding FAST protein interactions. Thus, we were interested to determine the potential utility of oversized fluorogens as highly selective markers of cell viability. To test this, we clicked N-(acid-PEG10)-N-bis(PEG10-azide) to ^PL^Blaze-alkyne, as it is both commercially available and exceptionally bulky, adding three PEG arms of 40 Å each. Despite its enormous size following conjugation, ^PL^Blaze-PEG retained most of the normal function of its parent fluorogen, exhibiting an apparent brightness just 2-fold below that of ^PL^Blaze using both iFAST2 and frFAST2 proteins (**Table 1**). Compared with molecules like PI, diffusion of similarly PEGylated fluorogens through the cell membrane would require pores of at least an order of magnitude greater size to allow this ∼90 Å molecule access to the cytosol (**Fig. 3a**).

**Fig. 3.**
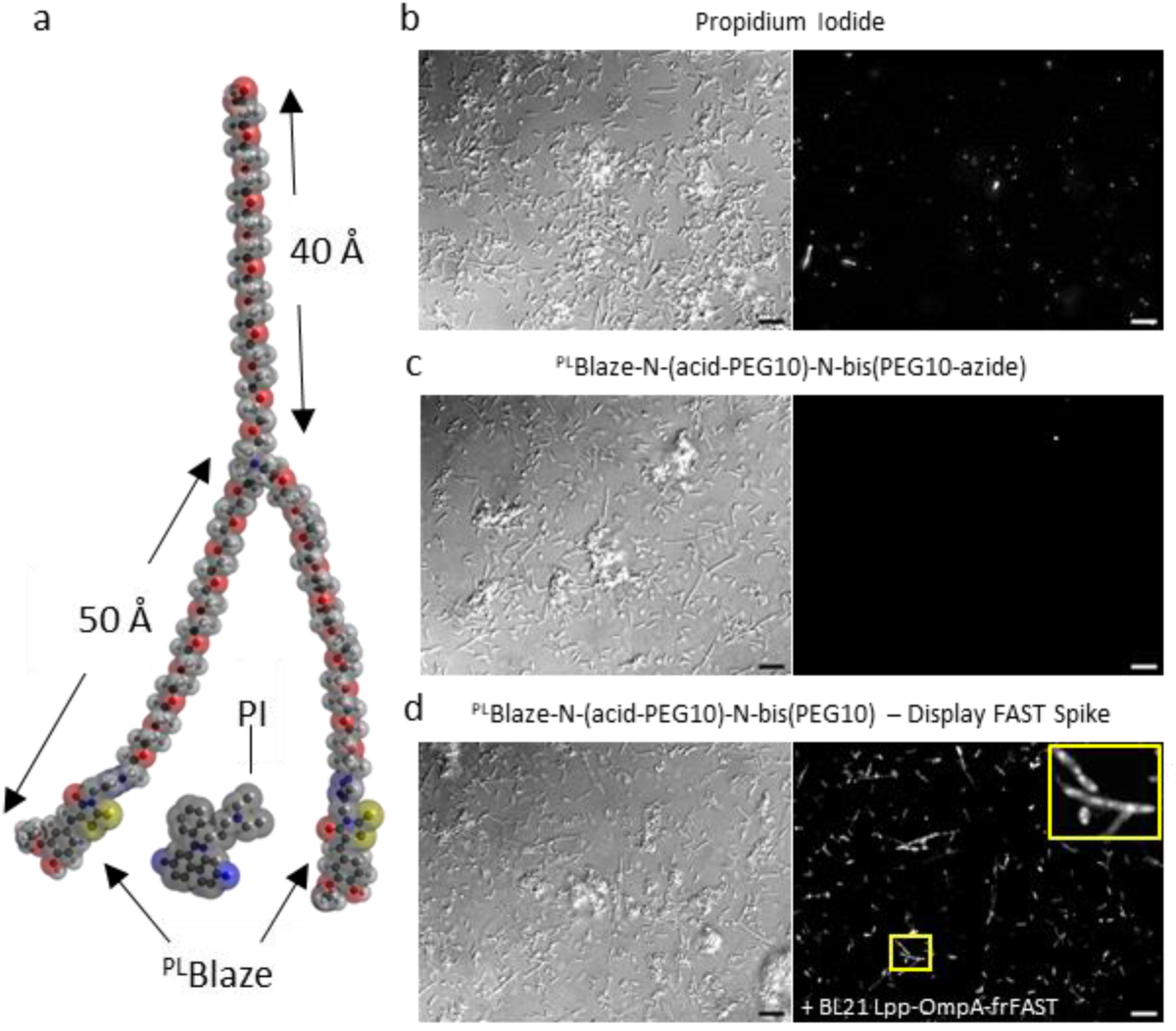
Click attachment of N-(acid-PEG10)-N-bis(PEG10) to ^PL^Blaze imparts stringent membrane impermeability. **a**, Model of ^PL^Blaze-PEG. Two arms span roughly 50 Å with fluorogens conjugated to them. A third arm spans a maximal distance of 40 Å. Propidium iodide is denoted with “PI” between two ^PL^Blaze fluorogens to demonstrate comparative scale. **b**, DIC image (*left*) and fluorescence maximum intensity projection image (*right*) of a mixed population of frFAST2-expressing *S. mutans* UA159 and *E. coli* BL21 in the presence of propidium iodide. **c**, DIC image (left) and fluorescence maximum intensity projection image (*right*) of a mixed population of frFAST2-expressing *S. mutans* UA159 and *E. coli* BL21 in the presence of ^PL^Blaze-PEG. **d**, DIC image (*left*) and fluorescence maximum intensity projection image (*right*) of a mixed population of frFAST2-expressing *S. mutans* UA159 and *E. coli* BL21 in the presence of ^PL^Blaze-PEG after spiking the culture with *E. coli* BL21 expressing frFAST on its outer membrane via an Lpp-OmpA fusion. Image intensity is matched to c. *Inset*, Zoom view of fluorescent BL21 *E. coli* with characteristic OmpA-FAST staining. Scale bars = 10 µm.

We conducted imaging experiments by combining constitutive frFAST2-expressing mid-logarithmic phase cultures of *S. mutans* and *E. coli* in fresh media containing either ^PL^Blaze-PEG or a molar equivalent of propidium iodide. Despite incubating the cells under stress-free growth conditions, PI labeling was apparent in a subpopulation of both organisms (**Fig. 3b**). However, the equivalent conditions resulted in virtually undetectable PEG-FAST staining, suggesting the PEGylated fluorogen is completely impermeable to both species (**Fig. 3c**). To exclude the possibility that negative staining was attributable to other factors inhibiting fluorescence, we next spiked these same cultures with a strain of *E. coli* BL21 expressing OmpA-frFAST. After adding this strain to the non-fluorescent mixed cultures, FAST fluorescence was easily detectable from extracellularly displayed FAST, indicating the PEGylated ^PL^Blaze in the medium was indeed fully functional (**Fig. 3d, Suppl. Fig. 3**).

We next sought to perform a similar analysis using an antimicrobial dose response curve. Chlorhexidine (CHX) is a broad spectrum antiseptic commonly used in many clinical settings ^29^. This molecule contains two chlorophenyl and biguanide groups flanking a methylene chain, resulting in a cationic nature that destabilizes negatively charged microbial membranes. Low CHX concentrations are bacteriostatic due to disruptions of osmotic equilibrium, whereas high concentrations are bactericidal, leading to lysis and coagulation ^30^. We performed a dilution series with CHX using frFAST2-expressing *E. coli* and observed a bactericidal effect at 0.002-0.006% CHX (**Fig 4a**). Differential interference contrast (DIC) images of these cells revealed predominantly intact organisms up to 0.006% CHX, which is close to the limit where viability dissipates in this assay. The highest exposure of CHX tested (0.02%) resulted in bright staining by PEGylated ^PL^Blaze, followed by a faint fluorescence at 0.006% and nearly undetectable levels thereafter using identical exposure settings. As expected, PI yielded bright fluorescence at the highest CHX dose, but significant fluorescence remained detectable down to 0.002% CHX, where cell viability is minimally affected. Only faint signals were observed at 0.0007% CHX, illustrating the dose-dependence required for PI staining (**Fig. 4b**).

**Fig. 4.**
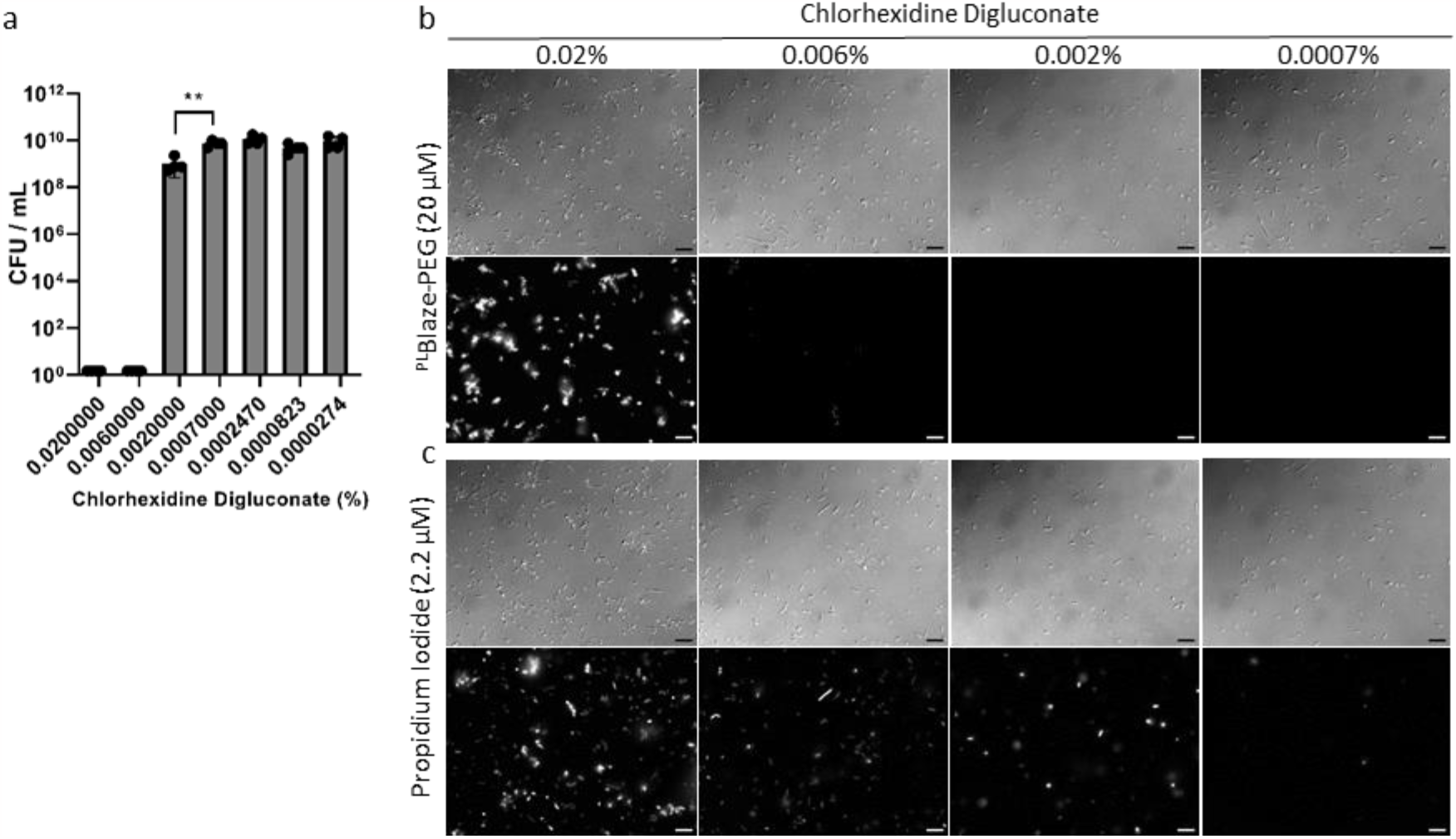
Performance of propidium iodide versus ^PL^Blaze-N-(acid-PEG10)-N-bis(PEG10) in the presence of chlorhexidine. **a**, Comparative colony forming units per mL of *E. coli* BL21 in the presence of the indicated concentration of chlorhexidine after 2 h incubation at room temperature. N = 4 ± SD. **, P = 0.001. **b**, *Top row*, Representative DIC images from three experiments examining *E. coli* BL21 in the presence of 20 µM ^PL^Blaze-PEG and the indicated chlorhexidine concentrations. *Bottom row*, ^PL^Blaze-PEG signal corresponding to the above DIC image. The scale of the signal intensity was equally matched for the four images. Scale bars = 10 µm. **c**, Representative images for identical experiments as described in **b**, except the cells were incubated in the presence of 2.2 µM propidium iodide.

Finally, we sought to further compare the performance of PI versus PEGylated fluorogens for viability measurements using *S. mutans* as a representative monoderm bacterium. Specifically, we were interested to determine whether stationary phase cultures labeled with PEG-FAST would exhibit similar fluorescence as we have routinely observed with PI, despite the cells being fully viable. Overnight stationary phase cultures were diluted with 1% sucrose, which is a standard protocol used to initiate *S. mutans* biofilm formation. In agreement with our previous observations, we detected robust PI staining of the cells, particularly within early-stage biofilm microcolonies (**Fig. 5)**. In contrast, samples labeled with PEGylated ^PL^Blaze yielded little to no detectable fluorescence unless 0.1% CHX was added to induce cell death. When considering the results of the ^PL^Blaze-PEG labeling experiments of both *E. coli* and *S. mutans*, we conclude that different levels of membrane permeability should be detectable microscopically by conjugating various membrane-impermeable molecules to Click-FAST fluorogens.

**Fig. 5.**
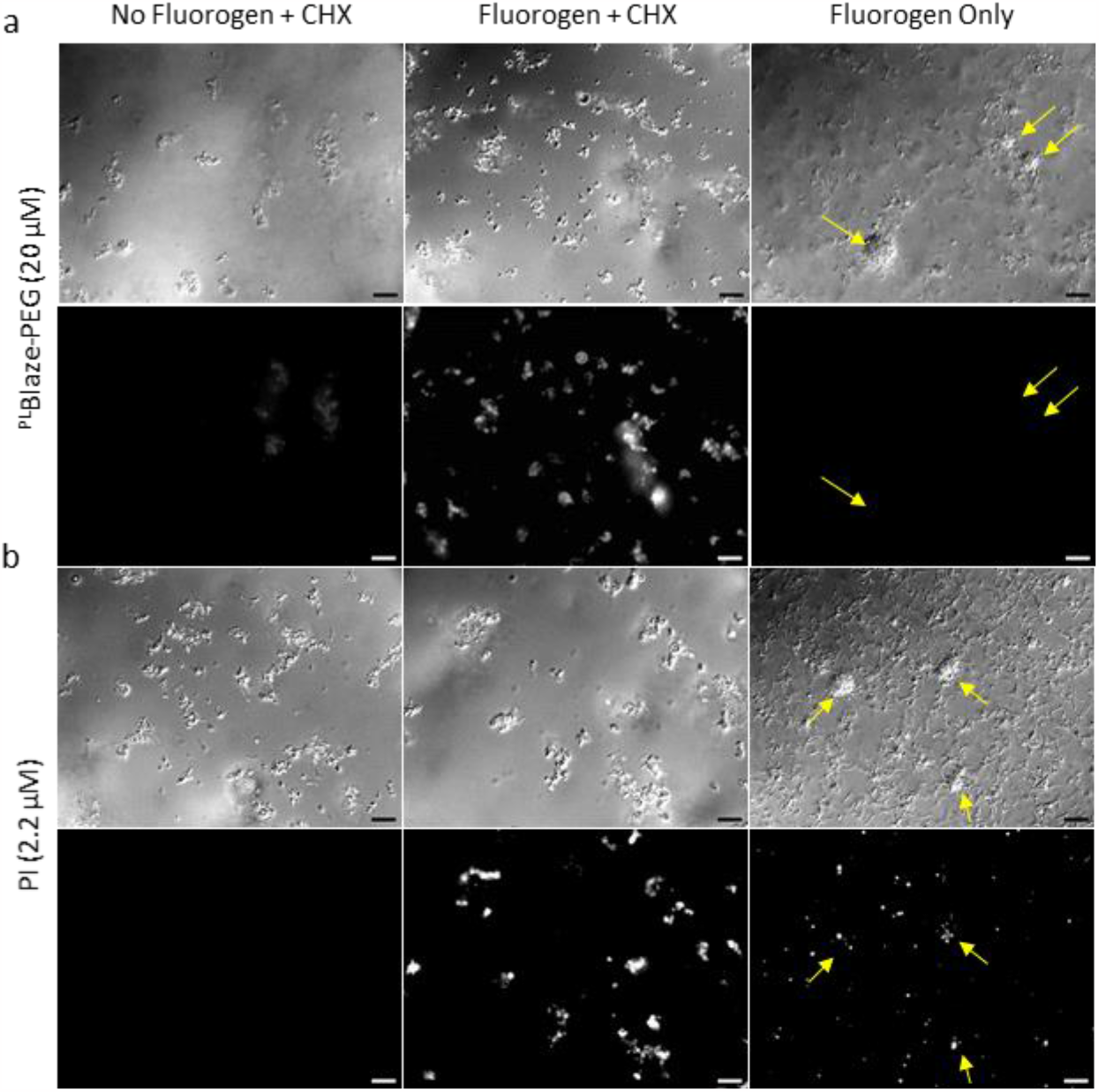
^PL^Blaze-N-(acid-PEG10)-N-bis(PEG10) exhibits improved performance over propidium iodide in stationary phase *S. mutans* UA159. **a**, *Top row*, Representative DIC images from three experiments examining stationary phase *S. mutans* UA159. *Top left*, No fluorogen with 0.1% chlorhexidine in the medium. *Top center*, 20 µM ^PL^Blaze-PEG and 0.1% chlorhexidine. *Top right*, 20 µM ^PL^Blaze-PEG in the medium. *Bottom row*, Maximum intensity projections of ^PL^Blaze-PEG signal corresponding to the above DIC image. The scale of the signal intensity was equally matched for the three images. Scale bars = 10 µm. **b**, Representative images for identical experiments as described in **a**, except the cells were incubated in the presence of 2.2 µM PI. Arrows indicate microcolonies indicative of early-stage biofilms.

## Discussion

Innovation and development around the FAST system is progressing at an impressive pace. New fluorogen and protein variants are consistently being reported to address an array of specialized needs. These include new color options, split protein reporters, biosensing applications, and optically-tunable or caged head groups. Our motivation was to deliver a fluorogen variant that would be relatively inexpensive, easily sourced, and modifiable by nearly any laboratory according to specific user-defined needs. The building blocks of the base fluorophore, ethylvanillin and rhodanine, can be typically purchased for $1 – $2 per gram, resulting in enormous yields of fluorogen for a minimal investment. Furthermore, the synthetic scheme (**Suppl. Fig. S1**) used in this study to create a clickable rhodanine (**Suppl. Fig. S2**) is a straight-forward process that could be replicated at most institutions with a chemistry department. These fluorogens can be used in normal FAST fluorescent imaging applications, but are also compatible with click chemistry should a user need to impart additional biological activities to the fluorogen.

Click chemistry is ideally suited to support user-defined modifications, as it simply requires one to mix a small number of reagents, all of which can be performed in dimethyl sulfoxide, the preferred solvent used to suspend FAST fluorogens. We envision this approach to be particularly powerful for investigators aiming to screen compounds that functionalize fluorogens with unique and selective properties. A major advantage of the FAST system is that no fluorescence is produced until the fluorogen is complexed with the FAST protein^7^. Thus, one could utilize Click-FAST in high-throughput library screens to probe for molecules that are transported to the cytosol of a desired cell type. Furthermore, chemical libraries developed via the diazotization methods described by Meng and coworkers ^31^ would be directly compatible with this approach. Collaborative developments within the FAST imaging community could then potentially produce lead molecule targets for new synthesis campaigns. Another outstanding need for FAST imaging technology is the development of more selective protein-fluorogen combinations. By combining high-throughput fluorogen conjugation with directed evolution of FAST proteins, it is conceivable that new cognate protein-ligand pairings could be developed to support more effective multiplex imaging applications.

We demonstrate in this work two use cases for clickable FAST fluorogens. The first involves the differential permeability for a particular cell type. We found that *S. mutans* exhibited a far greater propensity to import a benzoic acid modified fluorogen compared to *E. coli* (**Fig. 2a, c**). The physiological underpinnings of this phenomenon have yet to be determined, but the results illustrate how useful fluorogen modifications could be identified from click chemistry libraries to discover reagents imparting selective cell permeability or active/passive transport mechanisms needed for a given model system. Moreover, the successful implementation of N-(acid-PEG10)-N-bis(PEG10-azide) conjugation (**Fig. 3a**) demonstrates the exceptionally wide range of clickable molecules that could be sampled in library screens. PEG conjugation also has additional applications as a reagent for microscopic assessments of cell viability, which is typically achieved using PI as a measure of membrane integrity. Here, we illustrated a major limitation of PI staining for microscopic viability measurements by enumerating CFU counts vs. PI fluorescence imaging in the presence of chlorhexidine, a commonly used membrane-disruptive antiseptic (**Fig. 4**). This was further illustrated with PI staining of stationary-phase *S. mutans* cells (**Fig. 5**). Thus, these data should serve as a cautionary tale for employing PI to quantify viable cells in microscopy studies requiring live/dead cell metrics. However, PEGylated fluorogens offer tremendous potential to achieve these same goals with much greater precision.

In summary, the FAST imaging system is proving itself to be of outstanding versatility for anaerobic live cell imaging and is poised to revolutionize fluorescence microscopy studies of the human microbiome. A wide range of pathologies occur exclusively in hypoxic/anoxic conditions, and the FAST platform offers the research community an attractive method to study polymicrobial interactions and the host-microbe interface directly within this environment. With Click-FAST, a near limitless array of ligand modifications and bioactivities are possible, and the continued development of high-density click libraries will likely further expand these options in the future.

## Methods

### Chemical reagents

The following stocks were purchased for imaging, fluorogen synthesis and click assembly, and used without further purification: 4-Hydroxy-3-methoxycinnamaldehyde (458-36-6, Sigma), 3-ethoxy-4-hydroxybenzaldehyde (121-32-4, Sigma), N-(acid-PEG10)-N-bis(PEG10-azide) (2803119-06-2, Broadpharm), β-Lactose-PEG3-azide (246855-74-3, AFG Scientific), 4-azidobenzoic acid (6427-66-3, Sigma), propidium iodide powder (25535-16-4, Sigma), chlorhexidine digluconate solution (20% solution in water, 18472-51-0, Sigma).

### Alkyne-FAST ligand Synthesis

*3-(2-Propyn-1-yl)-2-thioxo-4-thiazolidinone*: Bis(carboxymethyl)trithiocarbonate (7.00 g, 31.0) mmol) and triethylamine (1.0 mL, 15.5 mmol) where dissolved in isopropanol (5 mL) and heated to 80 °C. Propargylamine was added and the reaction was stirred at 80 °C for 12 hours. After cooling, the solution was diluted with 100 mL ethyl acetate and the organic was washed five times with DI water (20 mL), twice with brine (20 mL), dried over sodium sulfate, and concentrated under vacuum to a dark red oil. The crude oil was further purified by column chromatography (1:9 EtOAc:Hexanes) yielding a brown solid of the title compound (0.36 g, 13.7% yield). ^1^H NMR (600 MHz, Chloroform-d) δ 4.74 (d, J = 2.5 Hz, 2H), 4.03 (s, 2H), 2.22 (t, J = 2.5 Hz, 1H).

*5-(3-ethoxy-4-hydroxybenzylidene)-3-(prop-2-yn-1-yl)-2-thioxothiazolidin-4-one* (^PL^**Blaze-Click):** 3-(prop-2-yn-1-yl)-2-thioxothiazolidin-4-one (114 mg, 0.67 mmol) and ethyl vanillin (111 mg, 0.67 mmol) were dissolved in isopropanol (3mL). Ammonium acetate (103 mg, 1.34 mmol) was added and the solution was heated to 80 °C overnight. After cooling, solids were isolated and recrystallized out of toluene to yield orange solids of the title compound (154 mg, 72% yield). ^1^H NMR (400 MHz, DMSO-d6) δ 8.63 (s, 1H), 7.74 (s, 1H), 7.23 – 7.16 (m, 2H), 7.03 (d, J = 8.6 Hz, 1H), 4.87 (d, J = 2.5 Hz, 2H), 4.22 (q, J = 7.0 Hz, 2H), 2.80 (t, J = 2.5 Hz, 1H), 1.43 (t, J = 7.0 Hz, 3H).

*5-(3-(4-hydroxy-3-methoxyphenyl)allylidene)-3-(prop-2-yn-1-yl)-2-thioxothiazolidin-4-one* (^TF^**Poppy-Click):** 3-(prop-2-yn-1-yl)-2-thioxothiazolidin-4-one (165 mg, 0.93 mmol) and 4-hydroxy-3-methoxy-cinnamaldehyde (160 mg, 0.93 mmol) were dissolved in isopropanol (3mL). Ammonium acetate (143 mg, 1.85 mmol) was added and the solution was heated to 80 °C overnight. After cooling, solids were isolated and recrystallized out of toluene to yield dark red solids of the title compound (140mg, 46% yield). ^1^H NMR (400 MHz, DMSO-d6) δ 9.89 – 9.66 (m, 1H), 7.57 (dd, J = 11.7, 0.8 Hz, 1H), 7.39 – 7.29 (m, 2H), 7.11 (dd, J = 8.3, 1.9 Hz, 1H), 6.96 (dd, J = 15.0, 11.7 Hz, 1H), 6.81 (d, J = 8.1 Hz, 1H), 4.74 (d, J = 2.5 Hz, 2H), 3.85 (s, 3H), 3.29 (t, J = 2.5 Hz, 1H), .

### Molecular biology and cell culture

Plasmids pET29a Lpp-OmpA-far-red FAST (frFAST), cytosolic far-red tandem FAST (frFAST2) and pDL287 cytosolic tandem far-red FAST2 (*ldh* promoter) where constructed by standard cloning methods. Fresh chemical transformations of *E. coli* BL21 or competence-stimulating peptide based transformations of *S. mutans* UA159 were conducted before each experiment. BL21 pET-frFAST was grown in Luria-Bertani medium with 50 µg/mL kanamycin and *S. mutans* pDL278-frFAST in Todd-Hewitt medium with 0.3% yeast extract and 1000 µg/mL spectinomycin.

### Click reactions

The following click reagents were purchased from Fischer scientific and used without further purification: L-ascorbic acid (AAA1561319), Tris[(1-benzyl-1H-1,2,3-triazol-4-yl)methyl]amine (TBTA, NC1355341) and copper (II) sulfate pentahydrate (AC197722500). Overnight room temperature reactions were performed in DMSO with 0.5 mM copper-TBTA, 0.5 mM fresh ascorbic acid, 5 mM alkyne fluorogen and 6 mM azide ligands.

### Fluorescence imaging

Oxygen sensitivity experiments were performed by diluting 6 uL of overnight S. mutans UA159 pDL278-frFAST2 into 1 mL pre-reduced Todd-Hewitt both with 0.3% yeast extract (THYE) with 12.5 uM working Coral fluorogen in a MatTek dish. The vessel was placed into a stage top environmental chamber (Tokai Hit STX) for wide-field imaging on an Olympus IX73 inverted microscope. After a one hour, room temperature incubation period in 0.1% oxygen, 5% CO_2_, antibiotics were administered through a side port to bring the working concentration of each reagent to: 50 µg/mL gentamycin, 800 µg/mL kanamycin, 25 µg/mL erythromycin, 50 µg/mL chloramphenicol and 2 µg/mL tetracycline. This was followed by three rounds of mixture with a perfusion, media exchange and drug delivery system (Tokai PMD-D35) and after a further one hour incubation, the liquid media was evacuated by the PMD. Fluorescence images were acquired every minute using a mCherry filter. The next oxygen level was then set and 45 min was allowed to pass before the next imaging round. Images were identically processed using ImageJ Fiji and reported as an integrated fluorescence density normalized to the initial time point for each oxygen concentration.

Dual strain click-FAST imaging was conducted using BL21 and *S. mutans* UA159 grown to mid-log stage in their respective media. Cells were harvested at room temperature via a two minute, 16,873 x g centrifugation. Cells were suspended for imaging at an OD_600_ = 0.25 in pre-reduced THYE - 1% sucrose media without antibiotics. Click-^PL^Blaze fluorogen (20 µM final concentration) were added to the combined culture media and wide field images were acquired after a 1.5-2 hr of 37°C, 0.1% oxygen – 5% CO_2_ equilibration.

Chlorhexidine experiments with *E. coli* BL21 cells utilized cells harboring pET29a frFAST2. These were grown to mid-log phase in LB with 50 µg/mL kanamycin at 250 RPM at 37°C. Cells were diluted to an OD_600_ = 0.25 in fresh LB without antibiotics. Chlorhexidine water solution was added to the diluted culture at concentrations serially diluted between 0.02 and 0.0002% to be incubated for 2 h at room temperature. Cultures also either contained ^PL^Blaze-N-(acid-PEG10)-N-bis(PEG10) or propidium iodide at working concentrations of 20 or 2.2 µM, respectively. Cell viability was assessed via serial spot dilutions on LB agar media with 50 µg/mL kanamycin. Images were acquired on an Olympus IX73 inverted microscope in 0.1% oxygen and 5% CO_2_. Fluorescence signal was acquired in identical fashion for all images using a mCherry filter. Final images were identically processed and scaled within their fluorophore grouping.

Additional experiments with *S. mutans* UA159 utilized cells harboring pDL278 frFAST2 expressed via the lactose dehydrogenase promoter. Overnight stationary phase cultures were grown in THYE broth with 1000 µg / mL spectinomycin. Samples were combined with sucrose to one percent final concentration and imaged via wide-field microscopy similarly to the *E. coli* experiments.

## Supporting information

Supplemental Material for Click Fast Paper

## Acknowledgments

Support for this work was provided by NIDCR/NIH awards DE028252 to J.L.M., DE029612 and DE029492 to J.K., DE029083 to C.S.P., and an OHSU School of Dentistry Exploratory Research Seed Grant to D.M.A.

## Author contributions

D.M.A. and J.L.M. conceived the study and wrote the manuscript. D.M.A. conducted experiments. M.G.L., S.S.P, M.J.K. and C.S.P. provided resources and synthesized the Poppy-alkyne and ^PL^Blaze ligands. All authors listed contributed to data interpretation, manuscript revision and final approval.

## Competing interests

The authors declare no competing interests

## Notes

### Competing Interest Statement

The authors have declared no competing interest.

