## Supplemental Material for Click Fast Paper for "Microbiome imaging goes à la carte: Incorporating click chemistry into the fluorescence-activating and absorption-shifting tag (FAST) imaging platform"

[1] Division of Biomaterial and Biomedical Sciences, Oregon Health & Science University, Portland, OR, USA
[2]Department of Molecular Microbiology and Immunology, Oregon Health & Science University, Portland, OR, USA
[†]Current Address: Pacific Northwest National Laboratory, Richland, WA, USA

### Supplemental methods

***Recombinant protein production.*** Hexahistadine-tagged recombinant iFAST2 and far-red FAST2 were purified from *E. coli* BL21 after a 3 h, 37°C, and 0.5 mM IPTG induction of the proteins from the pET29a vector. Standard procedures for metal affinity purification were used after lysis via sonication. Eluted samples were concentrated via centrifugal filters and applied to a Superdex 200 Increase 10/300 GL column (Cytiva) equilibrated with 25 mM Hepes, pH 7.0 and 150 mM NaCl. Peak fractions were collected for concentration assessment using $A_{280}$ measurements before immediate storage of aliquots at -80°C. One-time freeze-thawed samples were used for fluorescence measurements in gel filtration buffer using a BioTek Cytation 5 Cell Imaging Multimode Reader (Agilent). Graphs were constructed using Prism 9.5.1 with non-linear regression fitting and a one-site specific binding model.

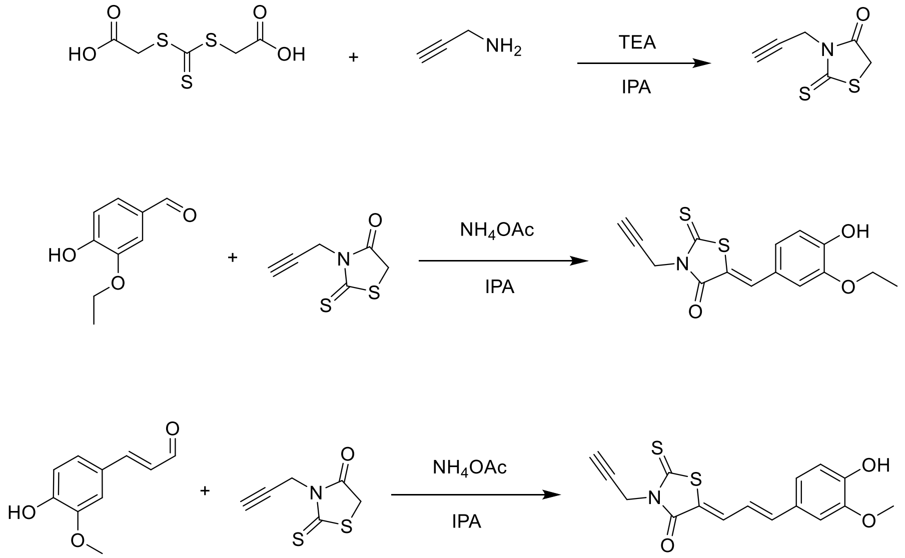


**Supplemental Fig. 1.** Synthetic schemes of *(top)* 3-(2-Propyn-1-yl)-2-thioxo-4-thiazolidinone, *(middle)* PLBlaze-Click and *(bottom)* TFPoppy-Click.

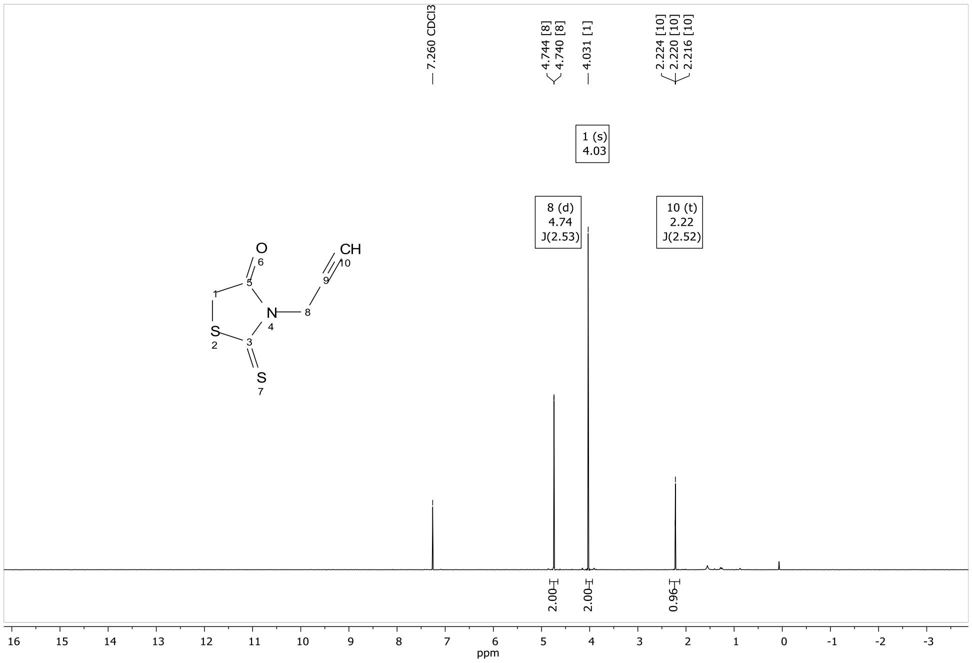


**Supplemental Fig. 2.** $^{1}H$ NMR spectra of 3-(2-Propyn-1-yl)-2-thioxo-4-thiazolidinone in $CDCl_3$.

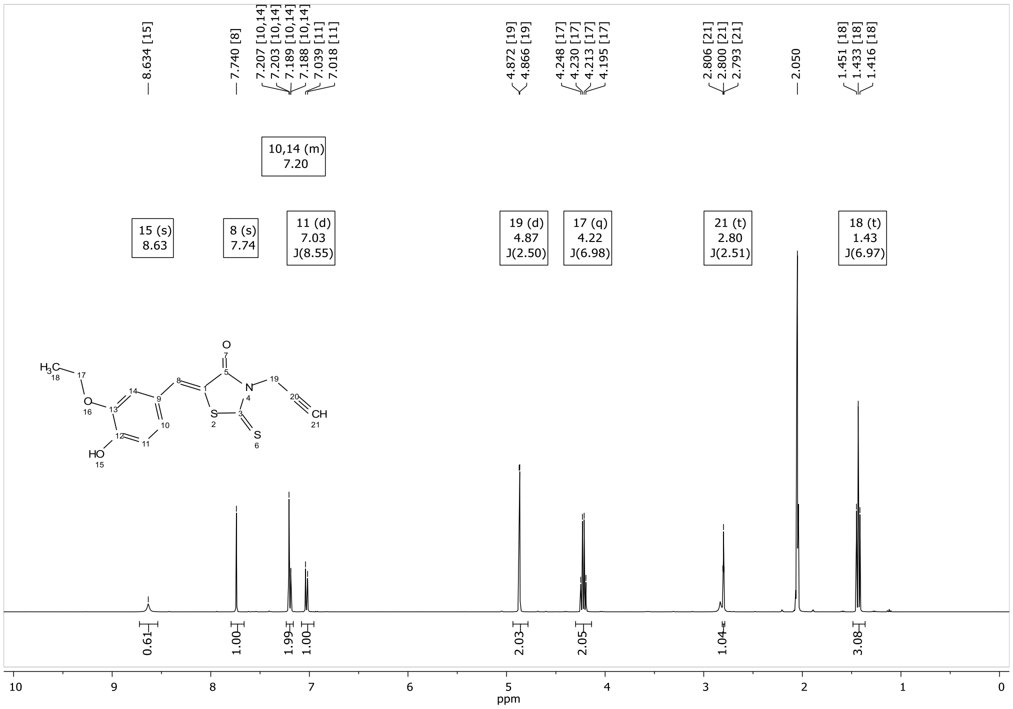


**Supplemental Fig. 3.** $^{1}H$ NMR spectra of 5-(3-ethoxy-4-hydroxybenzylidene)-3-(prop-2-yn-1-yl)-2-thioxothiazolidin-4-one ($^{PL}$**Blaze-Click**) in Acetone-$d_6$.

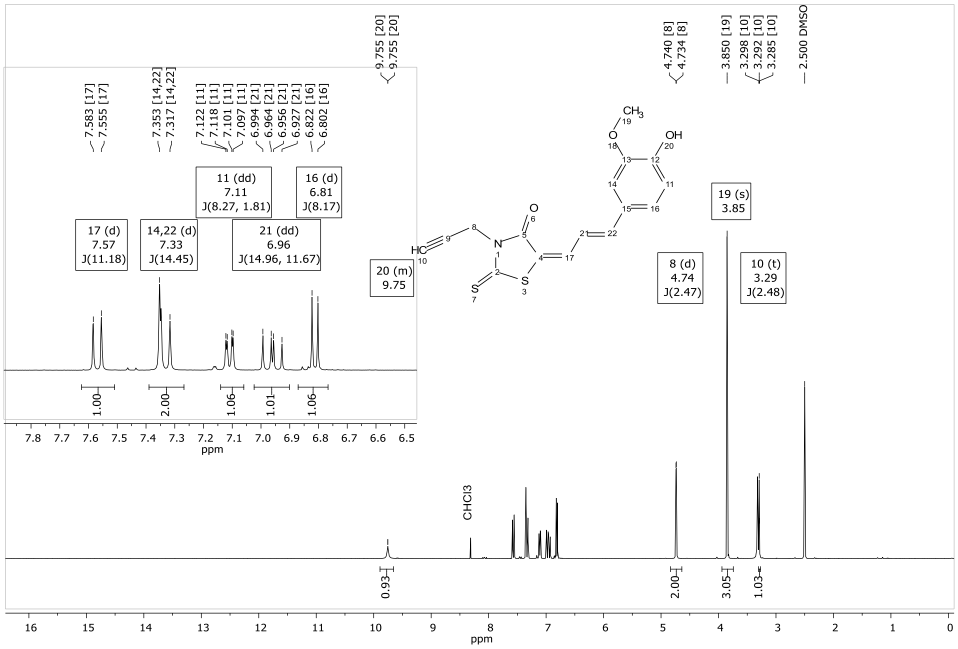


**Supplemental Fig. 4.** $^1$H NMR spectra of 5-(3-(4-hydroxy-3-methoxyphenyl)allylidene)-3-(prop-2-yn-1-yl)-2-thioxothiazolidin-4-one **($^{TF}$Poppy-Click)** in DMSO-$d_6$.

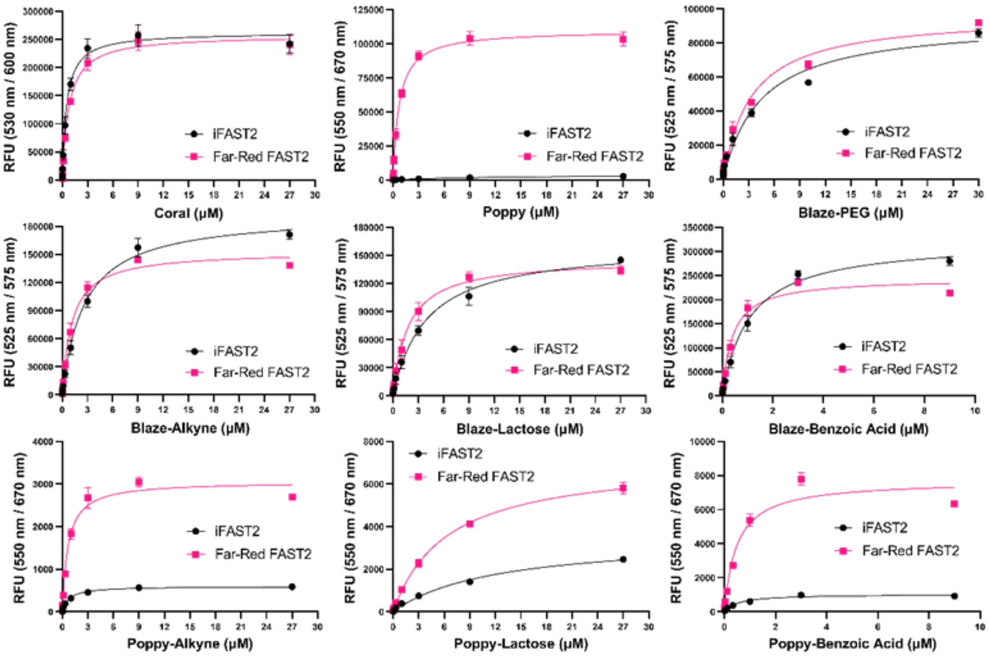


**Supplemental Fig. 5. FAST fluorogen fluorescence titrations.** iFAST2 or far-red FAST2 (500 nM) was combined with the indicated fluorogen for *in vitro* fluorescence measurements. The excitation / emission wavelengths are indicated for each series of measurements. N = 3 ± SD.

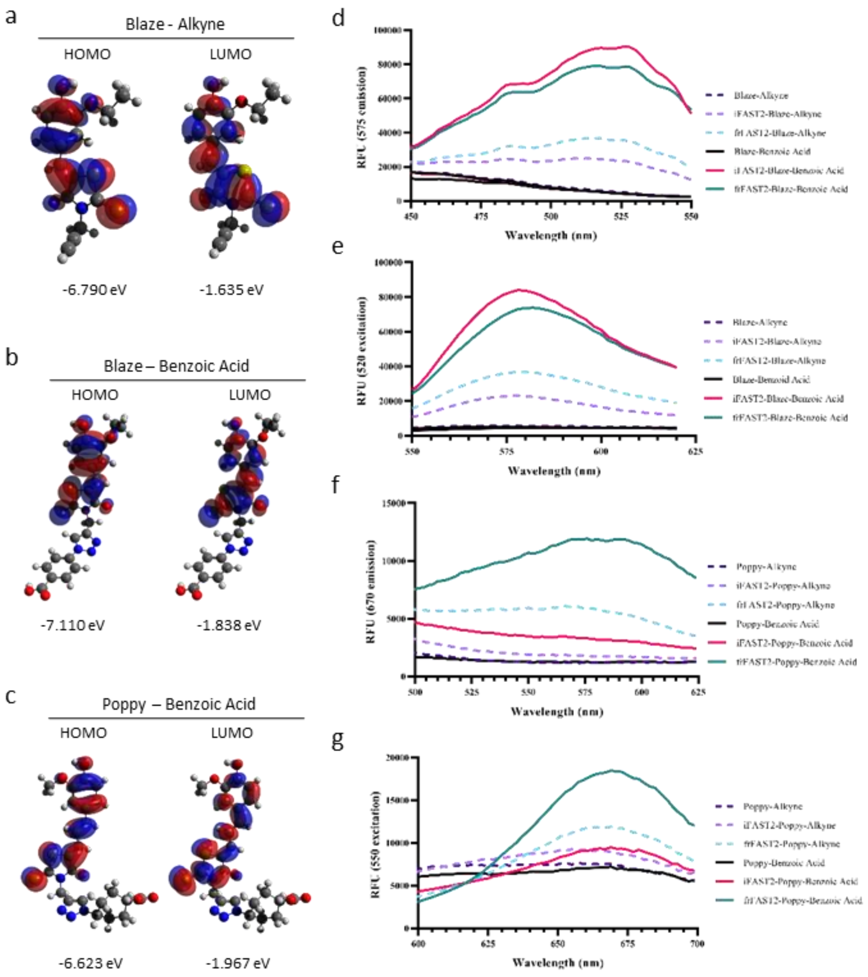


**Supplemental Fig. 6. Predicted electronic structures and spectral fluorescence. a**, HOMO and LUMO orbitals of a geometrically-optimized Blaze-alkyne fluoro. Orca 5.0.1[1] was used for all calculations using the following parameters: RIJK RI-PWPB95[2] D3BJ def2-TZVP def2/JK def2-TZVP/C TIGHTSCF. Orbital energy levels are shown below each model. **b-c**, Similar calculations for Blaze-benzoic acid and Poppy-benzoic acid, respectively. Models were generated from Avogadro version 1.2.0[3]. **d**, Excitation scan of Blaze-alkyne or Blaze-benzoic acid either in buffer or in the presence of iFAST2 or frFAST2. **e**, Corresponding emission scan to d. **f**, Excitation scan of Poppy-alkyne or Poppy-benzoic acid either in buffer or in the presence of iFAST2 or frFAST2. **e**, Corresponding emission scan to f.

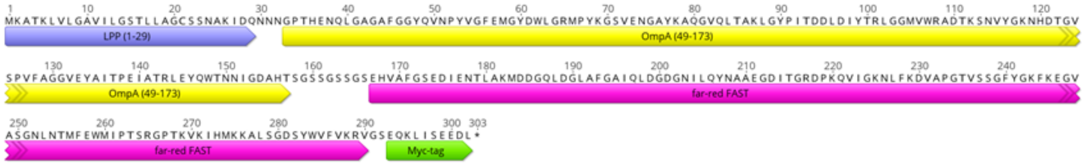


**Supplemental Fig. 7. Schematic of genetic fusion for surface-localized FAST in *E. coli*.** Amino acid sequence of Lpp-OmpA-far red FAST-myc. *Full sequence:*
MKATKLVLGAVILGSTLLAGCSSNAKIDQNNNGPTHENQLGAGAFGGYQVNPYVGFEMGYDW
LGRMPYKGSVENGAYKAQGVQLTAKLGYPITDDLDIYTRLGGMVWRADTKSNVYGKNHDTGV
SPVFAGGVEYAITPEIATRLEYQWTNNIGDAHTSGSSGSSGSEHVAFGSEDIENTLAKMDDGQLD
GLAFGAIQLDGDGNILQYNAAEGDITGRDPKQVIGKNLFKDVAPGTVSSGFYGKFKEGVASGNL
NTMFEWMIPTSRGPTKVKIHMKKALSGDSYWVFVKRVGSEQKLISEEDL.
